# Quantitative Analysis of Macrophage Uptake and Retention of Fluorescent Organosilica Nanoparticles

**DOI:** 10.1101/2023.06.12.544701

**Authors:** Hung-Chang Chou, Shih-Jiuan Chiu, Teh-Min Hu

## Abstract

This study investigates the uptake and retention of stable fluorescent organosilica nanoparticles by macrophages, which play a vital role in scavenging environmental nanoparticles and nanomedicine within the body. We used rhodamine 6G-loaded fluorescent organosilica nanoparticles (SiNP-R6G) synthesized from a thiol-functionalized organosilane precursor. Our primary objective was to establish a quantitative relationship between fluorescent measurements and nanoparticle tracking analysis, enabling the precise “counting” of nanoparticles taken up by macrophages under kinetic measurement conditions. Our kinetic study demonstrated a concentration-dependent, saturable internalization of nanoparticles in a model macrophage (RAW 264.7 cells), with a maximum uptake rate (*V*_max_) of 7.9 × 10^4^ nanoparticles per hour per cell. The estimated number concentration of nanoparticles for half-maximum uptake was approximately 0.8 trillion nanoparticles per milliliter, and a significant portion (∼80%) of internalized SiNP-R6G remained entrapped within the cells for 48 hours, indicating the sustained particle retention capacity of macrophages. These findings highlight the successful development of a methodology to accurately “count” the cellular uptake of nanoparticles in macrophages, providing valuable insights into the kinetics and retention capabilities of macrophages for nanoparticles.

## 1. Introduction

Nanomedicine expands the therapeutic horizon for new and existing drugs. However, despite numerous academic progress in the field, only a limited number of nanotechnology-based drugs have gained regulatory approval [1]. Improving the efficiency of nano-drug delivery is of utmost importance, as an analysis has shown that a mere 0.7% of nanoparticles successfully reach their intended tumor sites [2]. Macrophages play a vital role in the clearance and processing of nanoparticles, and their interactions with nanomaterials significantly affect the delivery efficiency to the target sites, thereby impacting therapeutic outcomes [3–5]. In this context, understanding the processes of nanoparticle internalization by macrophages becomes a critical aspect. Specifically, investigating the precise rate and extent of macrophage uptake and retention of nanoparticles can provide invaluable insights into optimizing nanoparticle delivery and enhancing overall therapeutic efficacy.

Numerous studies have focused on quantitatively analyzing the uptake and internalization of nanoparticles by macrophages, revealing insights into the intricate processes involved and their implications for nanomedicine. Specifically, these studies have demonstrated the crucial role played by the size, shape, and surface properties of nanoparticles in determining their uptake efficiency [6–17]. To gain a quantitative understanding of nanoparticle-cell interactions, researchers frequently employ various techniques, including confocal microscopy and flow cytometry, using fluorescent nanoparticles [18–27]. While these methods allow for relative comparisons based on fluorescence intensity between different treatment groups, they do not provide precise information regarding the “number” of particles internalized by cells. Alternatively, inductively coupled plasma (ICP)-based methods can be employed to measure the particle numbers within cells or tissues. This technique has been successfully applied to the analysis of inorganic nanoparticles like gold and silica nanoparticles [20, 22, 27–29], enabling the quantification of nanoparticle uptake. However, limited reports in the literature have explored the application of ICP-based methods for studies requiring continuous and intensive sampling for kinetic measurements. This could be attributed to the relatively limited accessibility of the specific instruments needed for the analysis. Accordingly, it is crucial to expand the available methodologies for quantitative cellular uptake studies.

Recently, there has been a resurgence of interest in modeling the cellular internalization of nanoparticles using kinetic models [30–33]. Such an approach offers quantitative insights into the interaction between cells and nanoparticles, specifically in relation to the uptake, transport, and retention processes [34]. Understanding these cellular kinetics is crucial for macrophages for two primary reasons. Firstly, macrophages play a significant role in the clearance of injected or inhaled nanoparticles [3, 4, 35]. Secondly, a new emerging role of macrophages involves their potential as carriers for delivering nanoparticles to pathological loci [36–40]. However, to address key questions regarding the rate and extent of these processes and enable dosage estimation and prediction, more precise measurements are required, specifically concerning particle number concentration, which governs the kinetic process [41].

Fluorescent nanoparticles have frequently been employed to trace the cellular internalization of nanoparticles [42]. Although versatile, this method is limited to providing semiquantitative information based on measured fluorescence intensity and is susceptible to potential dye leakage during the uptake process. To overcome these limitations, we propose a novel approach using organosilica nanoparticles loaded with a stable fluorescent dye, rhodamine 6G (R6G). Our method combines direct fluorescent measurement with nanoparticle tracking analysis to determine particle number concentrations during uptake and retention of nanoparticles in macrophages. To the best of our knowledge, similar approaches have not been published in the literature. In this paper, we present the measured kinetic data that enable the estimation of macrophage capacity for taking up and retaining organosilica nanoparticles. These results are corroborated by image analysis using confocal microscopy and transmission electron microscopy.

## 2. Materials and Methods

### 2.1 Chemicals

3-Mercaptopropyltrimethoxysilane (MPTMS, M0928) was purchased from Tokyo Chemical Industry Co., Ltd. Sodium bicarbonate was provided by Kanto Chemical Co., Inc. Dimethyl sulfoxide (DMSO) was acquired from J.T.Baker. All other chemicals were supplied by Sigma-Aldrich. Reagent-grade purified water was used in all experiments (18.2 MΩ·cm at 25 °C, ELGA LabWater, UK).

### 2.2 Preparation of Fluorescent Organosilica Nanoparticles

The rhodamine 6G-loaded organosilica nanoparticle (SiNP-R6G) was prepared using a modified nanoprecipitation method [43]. In brief, the organic phase consisted of DMSO with 3-mercaptopropyltrimethoxysilane (MPTMS, 200 mM), sodium nitrite (400 mM), diethylenetriaminepentaacetic acid (DTPA, 0.05 mM) and hydrochloric acid (500 mM). After reacting for 24 h at 25 °C, one milliliter of the organic phase was taken and injected into 10 mL of an aqueous solution of rhodamine 6G (0.08 mM) with ascorbic acid (80 mM). Then, after being kept in the dark for one hour, the nitric oxide-containing, S-nitrosothiol (SNO)-conjugated SiNP-R6G nanoparticles were collected and washed with purified water. Subsequently, a light-triggered decomposition of the nitric oxide (NO) moiety on SNO-SiNP-R6G was performed using a13W light source to obtain NO-free SiNP-R6G (refer to Scheme 1A). After 48 hours, the SiNP-R6G particles were thoroughly washed with purified water and stored in aluminum-foil wrapped containers for protection. To obtain a larger quantity of nanoparticles, repeat synthesis was conducted using additional organic phase solutions and repeated procedures, followed by concentrating the combined particle solutions through a centrifugation and redispersion procedure.

**Scheme 1.**
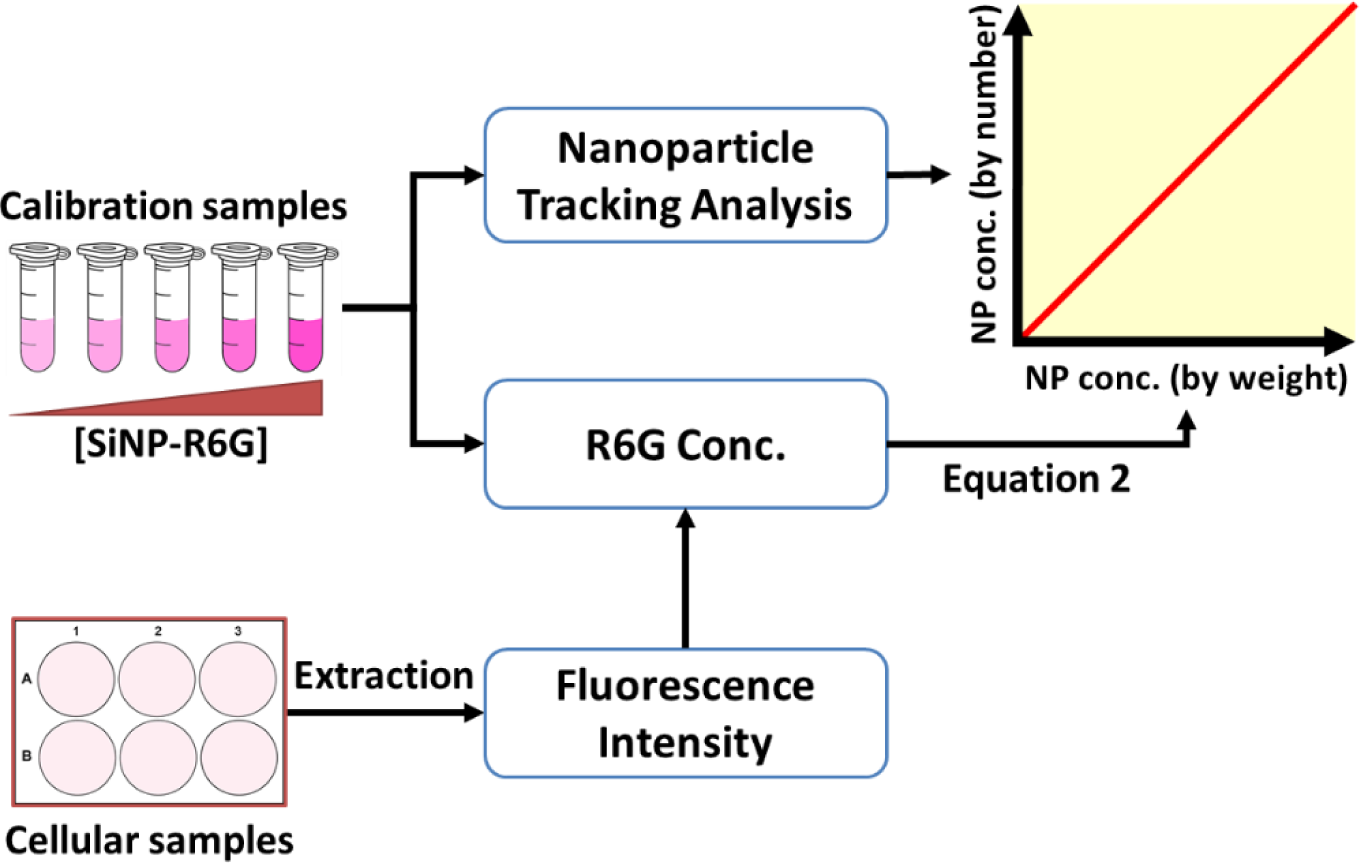
Schematic Representation of the Proposed Procedures for Measuring the Intracellular Number Concentrations of Fluorescent Nanoparticles. First, calibration solutions of SiNP-R6G nanoparticles were used to establish a quantitative relationship between the number concentrations and the mass concentrations of nanoparticles. The number concentrations of SiNP-R6G nanoparticles in 5 calibration samples were directly measured using nanoparticle tracking analysis (plotted on the y-axis). The mass concentrations of SiNP-R6G, plotted on the x-axis, were derived from R6G concentrations by applying Equation 2 to correct the loading content. For the cellular samples, intracellular fluorescence intensities were measured after extracting R6G from cell-internalized SiNP-R6G. Similarly, the measured R6G concentrations were converted to the mass concentrations of nanoparticles using Equation 2. Subsequently, the intracellular number concentrations of nanoparticles were estimated using the established quantitative relationship obtained from the calibration samples. (Refer to the main text for a detailed description of the procedures)

### 2.3 Characterization of SiNP-R6G

For the acquisition of transmission electron microscopy (TEM) images, colloidal particles were loaded onto a 300-mesh carbon-Formvar-coated copper grid (Electron Microscopy Sciences, Hatfield, USA) for 5 minutes. Excess solution was removed, and the grid was allowed to dry for 4 hours. TEM images were acquired using an accelerating voltage of 75 kV on a HT7700 TEM instrument (Hitachi, Japan).

Dynamic and electrophoretic light scattering techniques were utilized for the measurement of hydrodynamic particle sizes and zeta potentials, respectively (Zetasizer Nano ZS90, Malvern Panalytical Ltd., Worcestershire, UK). Adequately diluted samples (1 mL) were evaluated and the results were recorded.

To assess the kinetic stability of the nanoparticles, SiNP-R6G (final concentration equivalent to 1 mg/mL) was dispersed in various media, including water, 0.2 M acetic buffer pH 4.5, Dulbecco’s Modified Eagle Medium with or without fetal bovine serum. The hydrodynamic sizes, surface charges, and fluorescence intensity of the nanoparticles were measured at each time point. Fluorescence intensity was recorded using an excitation wavelength of 480 nm and an emission wavelength of 566 nm on a Spark 10M fluorescence reader (TECAN, Austria).

The rhodamine 6G content in the nanoparticles was determined using an alkaline extraction method. A stock particle solution was prepared by re-dispersing and diluting the as-prepared particles with water to achieve a final turbidity equivalent to absorbance = 1 (measured at 800 nm using a UV/Vis spectrometer, Spark 10M, TECAN, Austria). A diluted nanoparticle solution (20 μL) was dispersed in 980 μL of sodium hydroxide (1 M) and sonicated for 30 minutes. After centrifugation at maximum speed for 15 minutes, the transparent supernatant (200 μL) of the sample was neutralized with hydrochloric acid (40 μL, 5 M). The fluorescence intensity was measured using a fluorometer (Spark 10M, TECAN, Austria) at 480/566 nm for excitation and emission, respectively. A rhodamine 6G aqueous solution was used as the standard solution and subjected to the same procedure (5-750 nM in NaOH). On the other hand, SiNP-R6G (1 mL) was centrifuged at maximum speed for 15 minutes to remove the supernatant. The weight of the nanoparticles was obtained by hot-air drying at 60℃ overnight. The loading content of rhodamine 6G in SiNP-R6G was calculated using Equation 1.

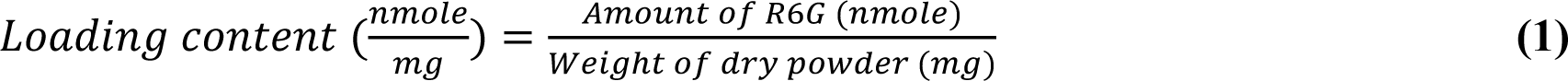

All experiments were independently performed in triplicate, and the results were recorded as mean ± standard deviation (SD).

### 2.4 The Relationship Between Nanoparticle Concentrations by Weight and Number: Estimation and Calibration

The relationship between estimated nanoparticle concentrations by weight and by number was established through two simultaneous measurements: fluorescence intensity of R6G and nanoparticle tracking analysis of SiNP (Scheme 1). To achieve calibration, freshly prepared SiNP-R6G stock particle dispersions with batch-based loading content were used. The stock solution was then serially diluted to prepare five nanoparticle solutions with corresponding R6G concentrations equivalent to 5, 10, 25, 50, 100 nM. The nanoparticle mass concentration (NP mass conc.; w/v) was calculated using Equation 2.

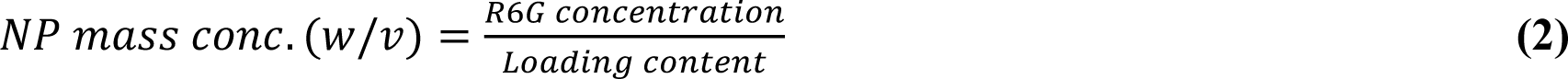

Simultaneously, the numerical concentrations of nanoparticles in each calibration solution were recorded using a nanoparticle tracking analyzer (NTA, NanoSight NS300, Malvern Panalytical Ltd., Worcestershire, UK), equipped with a 488-nm laser and an sCMOS camera. The Nanoparticle Tracking Analysis (NTA) software version 3.4.003 was employed. Each sample underwent three separate 60-second NTA measurements. Samples with a viscosity parameter of 0.9 cP (water) were carefully injected into the instrument chamber using a 1-ml syringe, ensuring no air bubbles were present. The NTA software used a detection threshold of 3 and automatically adjusted the maximum jump distance. The NTA charts displayed particle size distribution, with error bars indicating standard deviation.

Finally, the mass concentrations of particles were correlated with the number concentrations of particles using linear regression analysis. The deviation (%) between the estimated and observed number concentrations of particles was calculated by the following equation,

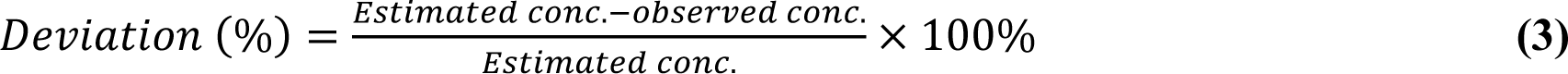

### 2.5 Cell Culture

RAW 264.7 macrophage cells were generously provided by Professor Jaw-Jou Kang’s group at National Yang Ming Chiao Tung University. The cells were cultured in Dulbecco’s Modified Eagle Medium (DMEM, D5648, Sigma-Aldrich) supplemented with 10% fetal bovine serum (FBS, 35-010-CV, Corning), 3.7 g/L sodium bicarbonate, 1 mM sodium pyruvate (25-000-CI, Corning), and a mixture of 100 IU/mL penicillin and 100 μg/mL streptomycin (30-002-CI, Corning), referred to as the complete culture medium. Cultivation was carried out in a humidified controlled atmosphere with 5% CO_2_, at 37 °C, with cells passaged every two or three days.

### 2.6 Cell Viability Assessment using MTT Assay

To evaluate cell viability, the MTT assay was performed. RAW 264.7 cells were seeded in Biofil^®^ microplates at a density of 5 × 10^5^ cells/well and incubated for 24 hours in a humidified atmosphere with 5% CO_2_ at 37 °C. The cells were then treated with either free R6G at concentrations of 0, 0.1, 0.25, 0.5, 0.75, and 1 μM, or SiNP-R6G at equivalent R6G concentrations of 0, 1, 2.5, 5, 10, and 15 μM (1 mL per well). After a 4-hour incubation, the drug-containing media were discarded, and the cells were washed twice with isotonic PBS. For acute toxicity assessment, cell viability assay was performed immediately. For subchronic toxicity evaluations, drug-free medium was added back to each well, and the cells were further incubated for 24 and 48 hours. After the designated time intervals, the cells were washed twice with isotonic PBS and subjected to viability assay. The cell viability assay involved the addition of 500 μL of MTT (0.5 mg/mL) to each well, followed by a 3-hour incubation. The formazan crystals formed were then dissolved by the addition of 500 μL of DMSO to each well. After shaking the microplate for 1 minute, the absorbance at 570 nm was measured using a spectrophotometer (Spark 10M, TECAN, Austria). The experiments were conducted four times, and the results were presented as mean ± SD (standard deviation).

### 2.7 Endocytosis and Exocytosis Experiments

For the endocytosis experiments, macrophages were seeded in Biofil^®^ 6-well cell culture plates (TCP-011-006) at a density of 5 × 10^5^ cells/well. The following day, the cell culture medium was removed, and the cells were exposed to various concentrations of R6G or SiNP-R6G for a specified duration of up to 4 hours, with each well representing one incubation duration. After the incubation period, the drug-containing media were discarded, and the cells were gently washed twice with iced PBS. Subsequently, 1 mL of 1 M NaOH (dissolving solution) was added into the microplates, followed by incubation for 30 minutes at 100 rpm using an orbital shaker (S101, Firstek Scientific, Taiwan). The solution was collected and processed as previously described for the measurement of rhodamine 6G content in the nanoparticles (Section 2.3).

For the exocytosis experiments, cells were seeded in the same manner as the endocytosis process, using Biofil^®^ 6-well cell culture plates (TCP-011-006) at a density of 5 × 10^5^ cells per well. After being seeded for 24 hours, the cells were exposed to R6G or SiNP-R6G at concentrations equivalent to 1 μM or 10 μM R6G for 4 hours. The dye-containing media were then removed, and the cells were gently washed twice with iced PBS. Subsequently, 1 mL of dye-free complete culture medium was added to the cells. At each designated time point, the culture media were discarded, and the cells were gently washed twice with iced PBS. The alkaline dissolving solution was added to the plates and incubated for 30 minutes at 100 rpm using an orbital shaker. The solution was collected and processed as previously mentioned.

### 2.8 Live Cell Imaging and Quantification

RAW 264.7 cells were cultured in bottom glass cell culture dishes (35 mm, Alpha Plus Scientific Corp., Taiwan) with complete medium at a density of 2.5 × 10^5^ cells per dish. The visualize the lysosomes, cells were stained with LysoTracker^TM^ Deep Red (1 μM, L12492, Thermo Fisher Scientific, USA) for 90 minutes at 37°C. Additionally, the cell nuclei were stained with Hoechst 33342 (10 μg/ml, H3570, Thermo Fisher Scientific, USA) for 20 minutes at 37°C. After staining, cells with clear morphology were selected based on observation in the bright field channel. The images were acquired under the following conditions: Hoechst 33342-excitation at 405 nm, emission at 460 − 490 nm (blue); R6G-excitation at 514 nm, emission at 530 – 590 nm (red); LysoTracker^TM^ Deep Red-excitation at 633 nm, emission at 650 – 800 nm (green); image size of 1,024 × 1,024 pixels in RGB format. The imaging was performed using a confocal laser scanning microscopy (CLSM, TCS SP5, Leica) with a 40X immersion objective and a live cell chamber system.

After treatment with SiNP-R6G (R6G concentration of 15 μM) or R6G (1 μM), cells were immediately and continuously observed using CLSM. After 4 hours, z-stack images were acquired with optical sections of 0.5 μm thickness and a size of 512 × 512 pixels. The z-stack images covered different focal depths (Z positions) within the cell. The fluorescent image with the minimum contour of the nucleus was designated as the zero position (Z0). Subsequently, the fluorophore-containing culture media were replaced with fresh medium. The following day, images were acquired to assess the retention of nanoparticles. The fluorescence intensity of each cell was quantified using ImageJ. Bright-field images were converted to 8-bit images, and the threshold was adjusted to the default settings. The black-and-white images were processed to fill any holes and analyzed as particles to determine the location of the cells. The mean fluorescence intensity of each cell was measured from the fluorescent images and compared using a paired t-test.

### 2.9 Transmission Electron Microscopy Imaging

RAW 264.7 cells were cultured in a chamber slide (154461PK, Thermo Scientific, USA) with complete culture medium at a density of 2.5 × 10^5^ cells per slide. The following day, the cell culture medium was removed, and the cells were exposed to a medium containing nanoparticles (R6G concentration equivalent to 15 μM) for 4 hours. Subsequently, the medium was discarded, and the cells were gently washed twice with low-temperature PBS. For fixation, the cells were initially treated with a fixative solution (2% paraformaldehyde and 2.5% glutaraldehyde in 0.1 M cacodylate) for 30 minutes. Afterward, the cells were rinsed three times with 0.1 M cacodylate (15 minutes per rinse) and then secondarily fixed with 1% OsO_4_ in 0.1 M cacodylate for 2 hours. Following another three rinses with cacodylate buffer (15 minutes per rinse), the cells were dehydrated using a series of alcohol concentrations (70% to 100%, increasing by 10% each time, and soaking for 15 minutes at each concentration). Subsequently, propylenoxide was added to the cells for three runs (10 minutes per run). The cells were then embedded in a propylenoxide/Epon mixture and vacuum-dried. The embedded samples were heated in an oven at 62°C for 2 to 3 days. Subsequently, the cells were sliced with a glass knife, resulting in sample sections with a thickness of approximately 70 − 80 nm. The images were acquired using a transmission electron microscope with an accelerating voltage of 75 kV (HT7700, Hitachi, Japan).

### 2.10 Data Analysis

The data were obtained from a minimum of three independent experiments and presented as mean ± SD. The differences among groups were analyzed using one-way analysis of variance (ANOVA) followed by Tukey’s multiple comparisons test. A p-value less than 0.05 was considered statistically significant.

## 3. Results and discussion

### 3.1 Preparation and Characterization of Fluorescent Nanoparticles: Synthesis and Analysis

In this study, we used rhodamine 6G (R6G)-loaded organosilica nanoparticles (SiNP-R6G) as the model fluorescent nanoparticles. The preparation of SiNP-R6G involved a three-step scheme (Figure 1A) using 3-mercaptopropyltrimethoxysilane (MPTMS) as the precursor [43]. In the first step, MPTMS was dissolved in DMSO and combined with a sodium nitrite aqueous solution (volume fraction of DMSO and aq. sodium nitrite = 8:0.5), forming the organic phase. Upon the addition of 5 M HCl to the organic phase, the solution immediately displayed the characteristic red color of primary S-nitrosothiols, resulting from the reaction between the sulfhydryl group of MPTMS and nitrous acid. The organic phase was kept in the dark for 24 hours, allowing the formation of polycondensed organosilica species [44]. In the second step, a portion of the organic phase was rapidly injected into an aqueous solution containing R6G. This procedure facilitated the entrapment of R6G within an organosilica matrix conjugated with S-nitrosothiols (SNO). In the final step, bright light irradiation was employed to remove the nitric oxide (NO) moiety from the nanoparticles. Our previous study has shown that this light treatment significantly enhances the fluorescence intensity of SiNP-R6G [43].

**Figure 1.**
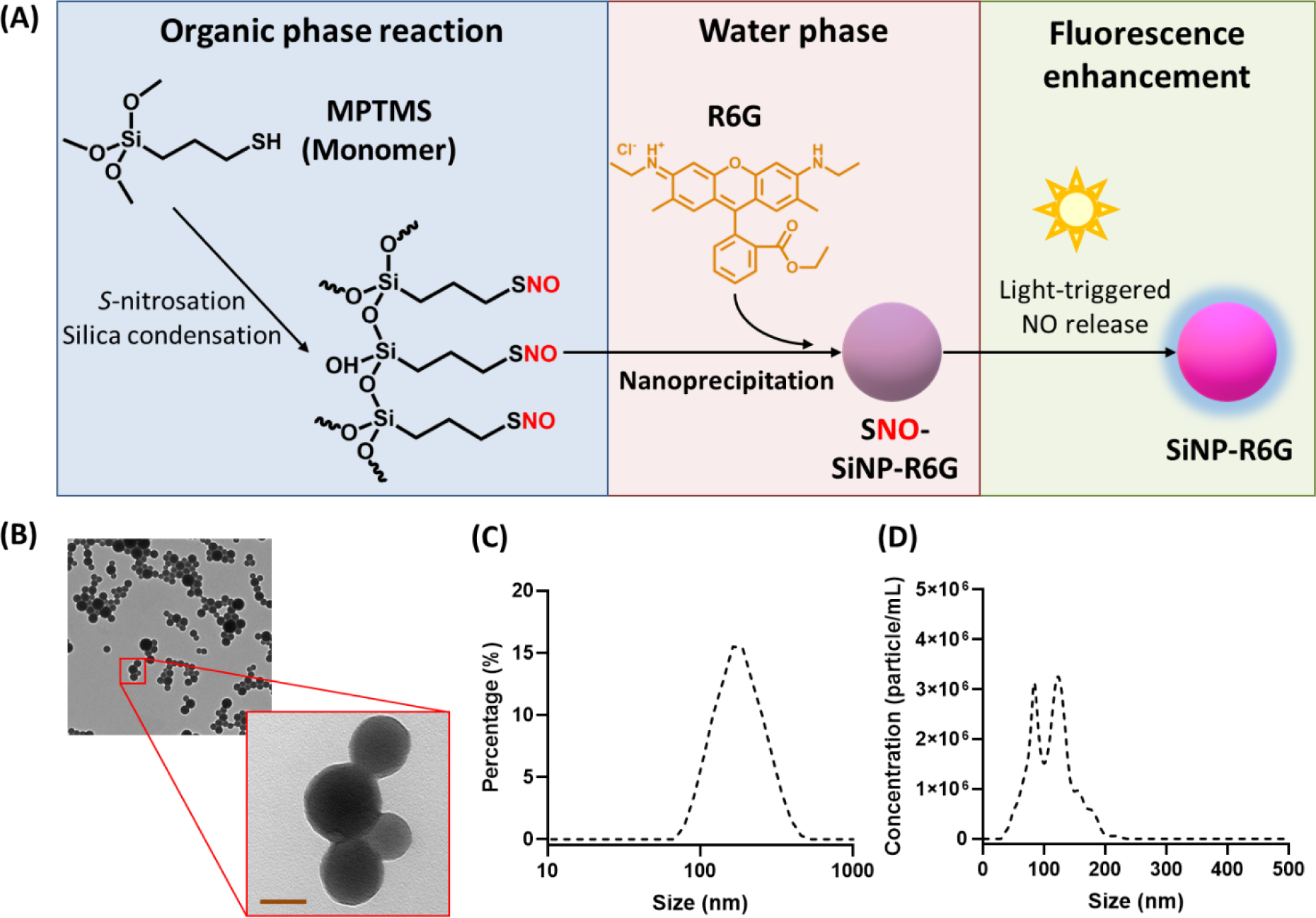
Preparation and Characterization of SiNP-R6G. (**A**) Schematic representation of the preparation process. MPTMS undergoes *S*-nitrosation and silica condensation, resulting in the formation of S-nitrosated (SNO) organosilica oligomers. These oligomers co-assemble with R6G into nanoparticles using a solvent-shifting (nanoprecipitation) scheme. After purification, the nanoparticles are exposed to bright light, which removes the NO moiety and enhances the fluorescence intensity of the final nanoparticles. (**B**) TEM images of the nanoparticles (scale bar = 50 nm). (**C**) Typical particle size distribution curve determined by dynamic light scattering (DLS). (**D**) Typical particle size distribution curved measured by a nanoparticle tracking analyzer (NTA).

The as-prepared SiNP-R6G nanoparticles underwent characterization to determine their particle sizes, zeta potential, loading content, and stability. The TEM images revealed solid spheres with a smooth surface (Figure 1B). The hydrodynamic sizes were determined to be161.0 ± 12.1 nm (by dynamic light scattering; DLS) and 124.0 ± 15.6 nm (by nanoparticle tracking analysis; NTA) (Figure 1C & 1D). The zeta potential of SiNP-R6G in water was found to be −60.7 ± 5.5 mV (n = 3). The encapsulated amount of R6G in SiNO-R6G was determined to be 37.0 ± 17.4 nmole per mg of nanoparticles (from 6 batches).

The colloidal stability of nanoparticles in cell culture media can be influenced by various components present, such as salts, amino acids and proteins [45]. Additionally, the colloidal stability profile may undergo alterations within the intraceullar environment, particularly in the acidic conditions of endosomes/lysosomes. Therefore, before using the fluorescent nanoparticles for cellular uptake studies, it is crucial to ensure their stability in the relevant media for the study system [18]. To assess the colloidal stability of SiNP-R6G nanoparticles, four different media were tested, including water, acetate buffer (pH 4.5), DMEM, and DMEM supplemented with 10% FBS at various time points up to 48 hours. Figure 2 illustrates the results obtained for the measurements of hydrodynamic size and zeta potential. Overall, the data demonstrate that the size and surface charge of the nanoparticles remain unchanged over time in all tested media. The observed variatoins in zeta potential among the different solutions indicate the presence of electrolytes and/or proteins in the media, beyond just water. SiNP-R6G exhibits a slighly larger size and less negative surface charge in FBS-DMEM compared to DMEM alone, suggesting interaction between serum proteins and the nanparticles. Importantly, the colloidal stability was confirmed in the culture medium, both with and without FBS, as well as in the acidic buffer mimicking the pH of intraceullar vesichles. The accompanying photoimages reveal no evidence of particle aggregation or sedimentation in the media relevant to the cell uptake experiment.

**Figure 2.**
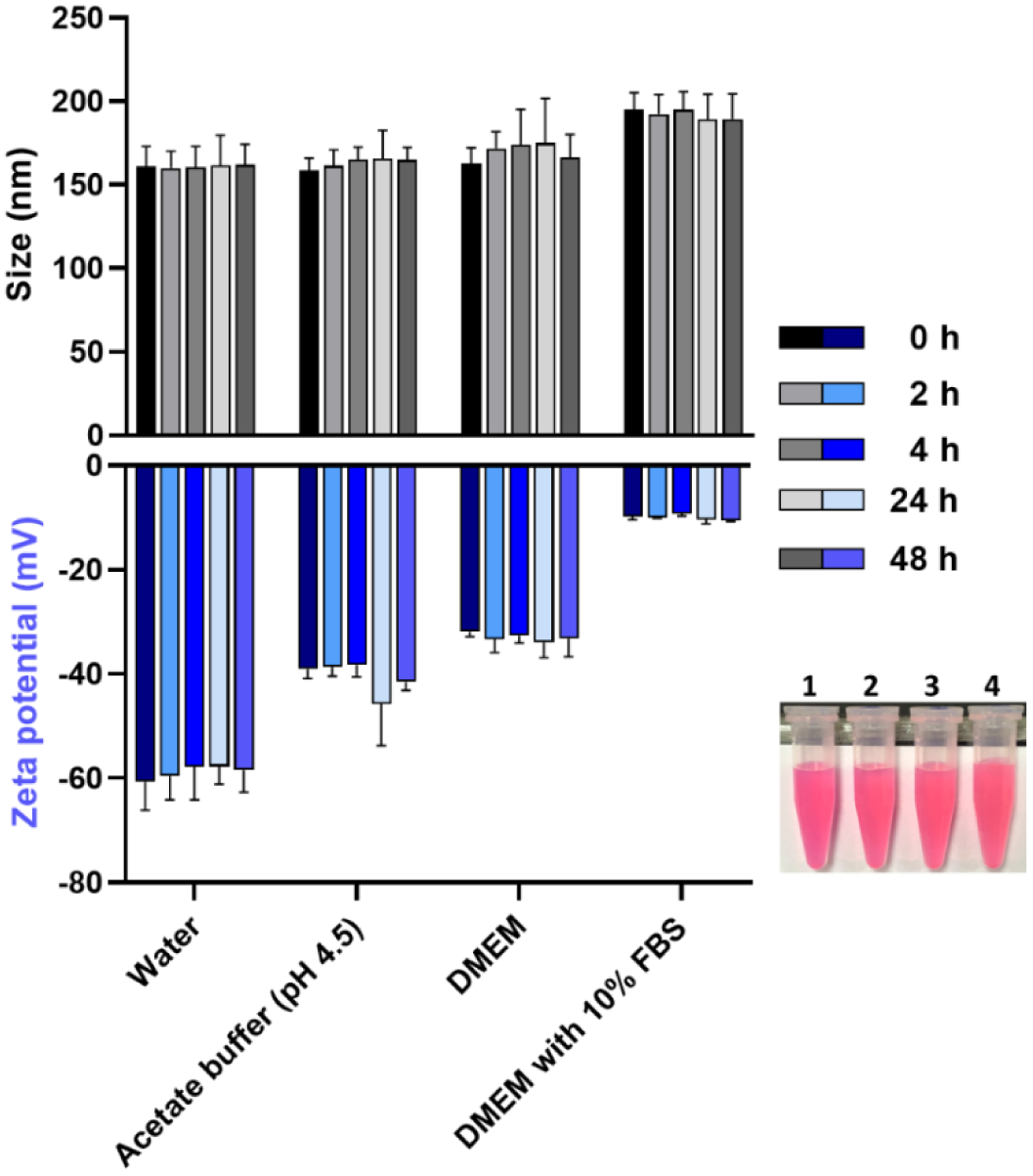
Stability of SiNP-R6G in Different Media. The hydrodynamic particle sizes and zeta potentials of SiNP-R6G remain constant over time when dispersed in various media. Data are presented as mean ± SD, n = 3. The photo image illustrates homogenous dispersion without an indication of particle sedimentation at 48 hour in the following conditions: (1) water, (2) acetate buffer (pH 4.5, 0.2 M), (3) DMEM, and (4) DMEM with 10% FBS.

The fluorescent stability of SiNP-R6G was aslo examined and confirmed. Figure 3 presents the fluorescnent intensity of SiNP-R6G in various media after a 48-hour incubation period. The accompanying photoimage displays the dispersion of particles (in FBS-DMEM) before and after centrifugation. The fluorescence intensity of the dispersions remains unchanged after suspension in the medium for 48 hours. Additionally, the supernatant, obtained after removing the nanoparticles via centrifugation, appears colorless and exhibits almost undetectable fluorescence. These results indicate that R6G is securely loaded within the nanoparticles without significant leakage over an extended period.

**Figure 3.**
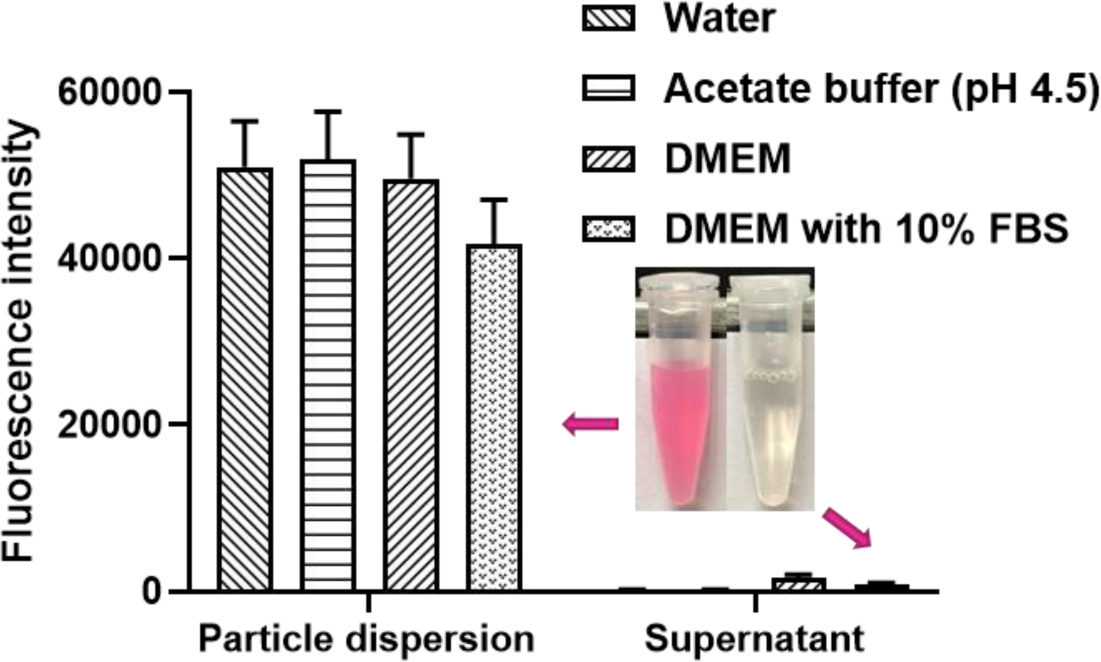
Stable Loading of R6G in SiNP-R6G Nanoparticles. SiNP-R6G nanoparticles were dispersed in different media to examine the stability of R6G loading. The fluorescence intensity of the particle dispersions was directly measured. After 48 hours, all dispersions were subjected to centrifugation, and the fluorescence intensity of the supernatant was measured. Data are presented as mean ± SD, n = 3. The photo image displays the dispersion of nanoparticles in the FBS-containing DMEM (left) and the corresponding supernatant (right) obtained from the same particle dispersion.

To use SiNP-R6G as a nanoparticle tracker for cellular uptake studies, it is essential to ensure that the nanoparticles do not adversely affect cell viability at the concentrations employed. RAW264.7 cells were subjected to incubation with SiNP-R6G at various equivalent R6G concentrations ranging from 1 μM to 15 μM. The MTT assay was used to assess cell viability at 4, 24, and 48 hours. The results indicate that cell viability remains unchanged regardless of the concentrations or treatment duration (Figure 4A). In contrast, free R6G exhibits concentration- and time-dependent cytotoxicity at lower concentrations (0.1 to 1 μM) (Figure 4B). These findings are consistent with the observation that the toxic R6G molecule is securely encapsulated within SiNP-R6G, leading to minimal release of free R6G.

**Figure 4.**
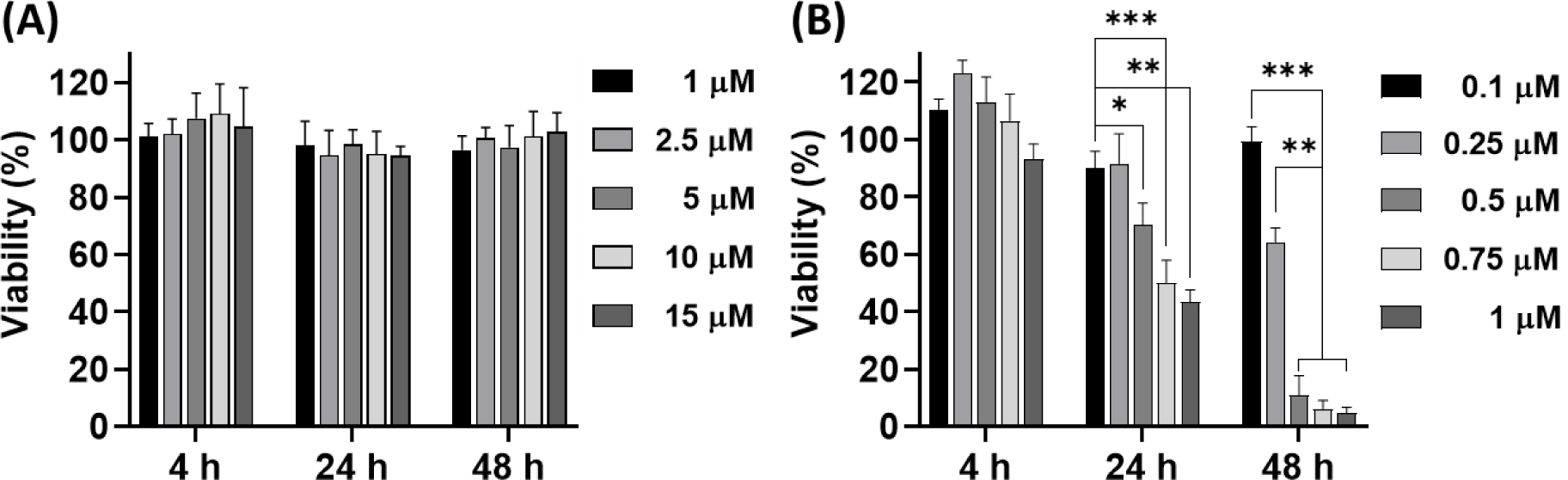
Viability of Macrophages Treated with SiNP-R6G and Free R6G. (A) Viability of macrophages treated with various concentrations of SiNP-R6G at different incubation times. The concentrations of SiNP-R6G are expressed as R6G equivalent concentrations. (B) Viability of macrophages treated with free R6G at different concentrations and incubation times.. Data are presented as mean ± SD, n = 4. * *p* < 0.05, ** *p* < 0.01, *** *p* < 0.001.

### 3.2 Conversion of Fluorescence Intensity to Nanoparticle Mass Concentration: The Nanoparticle “Counting” Process and its Relationship to Particle Number

In this paper, we propose a method to quantify the number of nanoparticles taken up by macrophages. The method combines two technologies: fluorometric determination and nanoparticle tracking analysis (NTA). We utilized SiNP-R6G to enable fluorometric tracking of the nanoparticles internalized by cells, assuming that each nanoparticle has a homogeneous and stable loading of R6G. Consequently, in the cellular uptake study, a simple extraction step can be employed to extract the fluorescent molecules internalized with nanoparticles in cells. To convert the fluorescence intensity into the exact number of nanoparticles, it is necessary to establish a quantitative relationship between the fluorometric measurement and the number concentration of nanoparticles in advance. The procedure is exemplified in Scheme 1.

To begin, six batches of SiNP-R6G were freshly prepared, purified, and re-dispersed in water to form stock SiNP-R6G solutions. The R6G concentration of the stock nanoparticle solution was determined by extracting R6G from SiNP-R6G using an alkaline solution. The concentrations of the extracted R6G were then estimated using a fluorometric standard curve of R6G (see Method section for details). Five calibration samples were obtained through serial dilutions of the stock solution. The number concentrations of the calibration samples were determined separately using NTA and plotted on the y-axis against the mass concentrations (weight-by-volume), which were estimated by normalizing R6G concentrations to the loading content of R6G (according to Equation 2) for each batch, on the x-axis.

In the cellular internalization experiments, the cellular samples were treated with alkaline extraction, followed by fluorometric determination of R6G concentrations. The intracellular R6G concentrations were then converted to the mass concentrations of nanoparticles, following the same procedures as described for the calibration samples. Finally, the intracellular number concentrations of nanoparticles were estimated using the established quantitative relationship between number and mass concentrations of nanoparticles.

Figure 5 depicts the calibration results obtained from 6 batches of SiNP-R6G. The number concentrations of particles exhibit a strong correlation with the mass concentrations of particles, displaying good linearity (R^2^ = 0.9697). The fitted linear equation is as follows:

**Figure 5.**
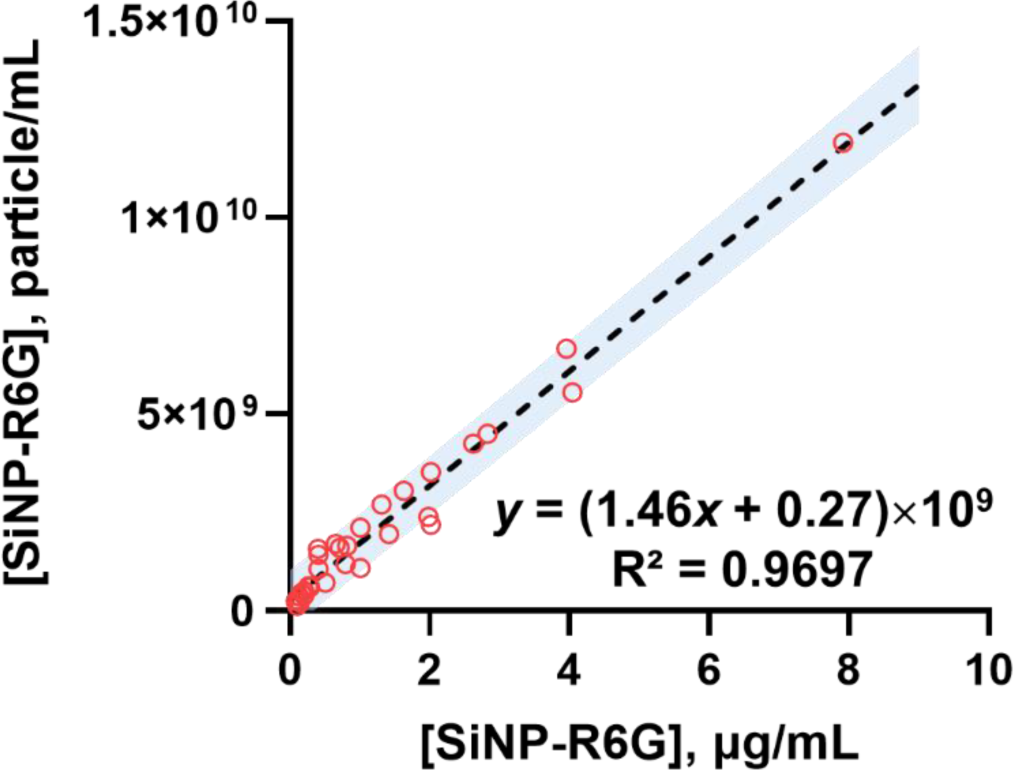
Relationship between the Estimated Mass Concentrations of Nanoparticles (*x*-axis) and the Number Concentrations of Nanoparticles Determined by NTA Measurements (*y*-axis). SiNP-R6G nanoparticles were freshly prepared, and the loading content (expressed as nmole of R6G in mg of nanoparticles) was determined. Five calibration solutions of SiNP-R6G were obtained through serial dilutions of the stock solution for NTA measurements. The concentration of nanoparticles (in μg/mL) on the *x*-axis was estimated using Equation 2, with the loading content serving as the batch-specific normalization factor. The *y*-axis represents the absolute number concentrations of nanoparticles obtained directly from NTA measurements. Refer to Scheme 1 and the main text for detailed procedures. The colored symbols correspond to independent experiments conducted on 6 batches of nanoparticles. The dashed line represents the regression line fitted to all data points, and the shaded area represents the 90% prediction interval.

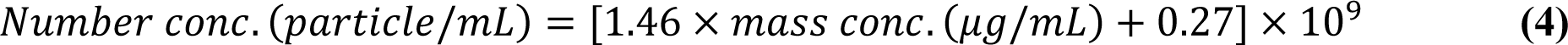

The estimated detection limit is 9.9 × 10^8^/mL [= 3.3 × standard deviation of the regression line/slope]. It should be noted that for each batch, the nanoparticle concentration range varies due to the batch-to-batch variation in R6G loading (since R6G concentrations were fixed in each batch). Furthermore, the dispersion of data points around the fitted line indicates the variability in the NTA analysis. However, the majority of the data points fall within the 90% prediction band. To validate the precision and accuracy of the proposed approach in estimating the number concentration of particles, a separate set of particle solutions containing three levels (low, intermediate, and high) of SiNP-R6G was employed (see Table 1 for a summary of the results). The coefficient of variation (CV%) is 32 % for the low-concentration sample and less than 6% for the samples with higher concentrations. The deviation between the estimated and observed number concentrations of nanoparticles for all levels falls within 20%. Additionally, the fluorescence readings remained largely unaffected when blank alkaline extracts of macrophages were added to free R6G solutions, suggesting that the cellular matrix does not interfere with the fluorometric determination of R6G. Consequently, the established quantitative relationship can be effectively used to estimate the number of nanoparticles internalized by macrophages.

**Table 1.** Precision and Accuracy of the Fluorometric Method for Predicting Particle Number

| **R6G conc.**<br>**(nM) <sup>a</sup>** | **µg of NP**<br>**per mL <sup>b</sup>** | **Estimated number**<br>**of NP per mL <sup>c</sup>** | **Observed number of**<br>**NP per mL <sup>d</sup>** | **Deviation**<br>**(%)** |
| --- | --- | --- | --- | --- |
| **20** | 0.36 | $0.79 \times 10^9$ | $0.82 \pm 0.27 \times 10^9$<br>(32%) | -3.8 |
| **75** | 1.34 | $2.23 \times 10^9$ | $1.83 \pm 0.11 \times 10^9$<br>(6.0%) | 17.9 |
| **150** | 2.68 | $4.19 \times 10^9$ | $4.02 \pm 0.18 \times 10^9$<br>(4.5%) | 4.1 |

- a. The R6G concentrations in three testing solutions represented low, intermediate, and high concentrations of SiNP-R6G nanoparticles. - b. The mass concentrations of nanoparticles were estimated using Equation 2. - c. The estimated number concentrations of nanoparticles were calculated using Equation 4. - d. The observed particle number concentrations were measured directly by NTA (mean ± SD, coefficient of variation (CV%) of 3 measurements).

### 3.3 Kinetics of SiNP-R6G Uptake in RAW264.7 Cells

The proposed fluorometric measurement with NTA calibration was used to study the kinetics of SiNP-R6G cellular internalization in the RAW264.7 macrophages. Initially, the uptake kinetics of free R6G were compared with those of SiNP-R6G at an equivalent R6G concentration (1 μM). The results indicate that the cells continued to internalize SiNP-R6G at a nearly constant rate, resulting in an uptake amount of approximately 13 ± 5% of the added amount after 4 hours. In contrast, the cellular uptake of free R6G rapidly reached a steady-state level of around 20% of the added amount within less than 30 minutes (Figure 6A).

**Figure 6.**
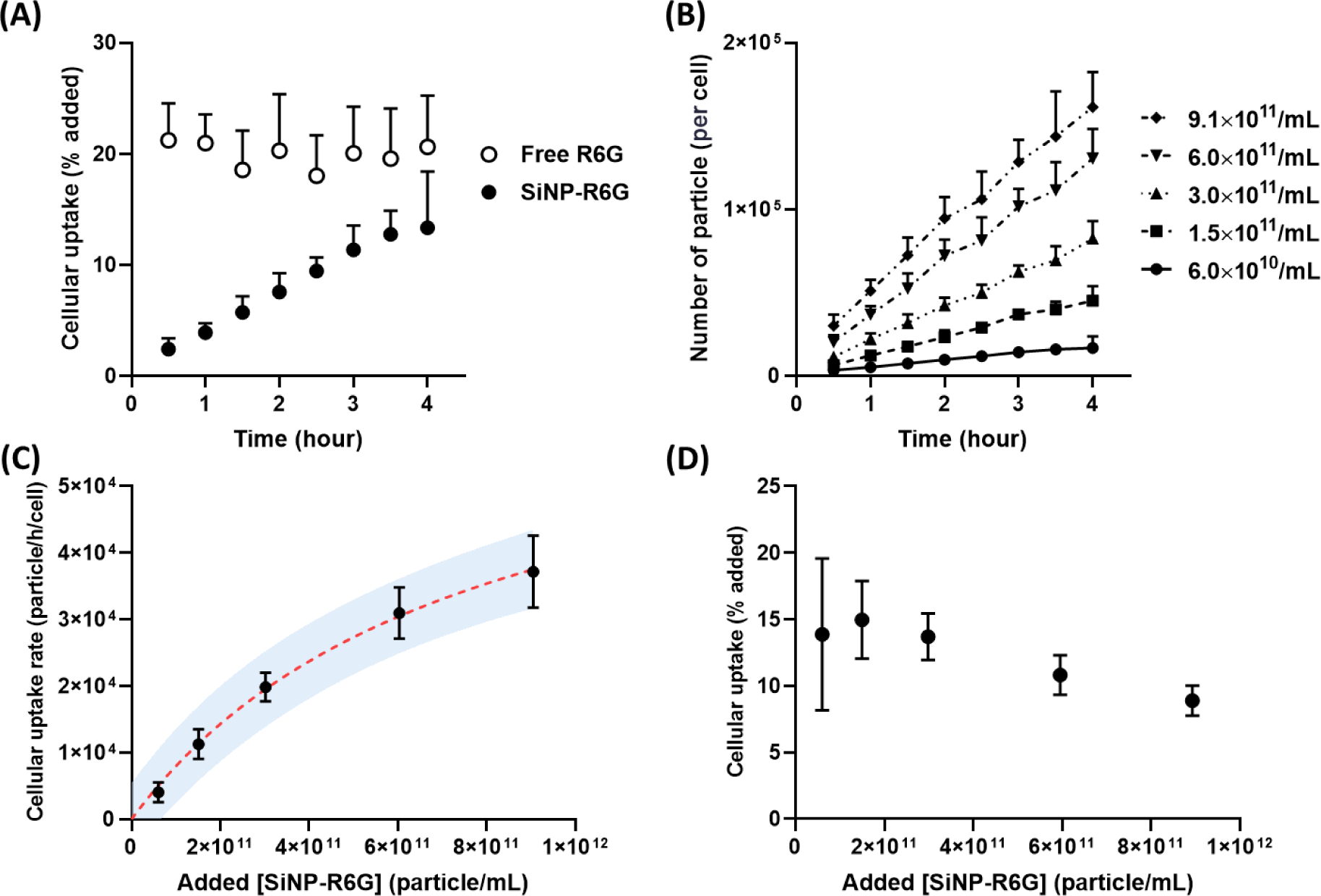
Uptake Kinetics of SiNP-R6G in Macrophages. (**A**) Comparison of uptake kinetics between free R6G and SiNP-R6G at an equal feeding concentration of R6G (1 μM). (**B**) The kinetic profiles illustrating the uptake of SiNP-R6G by macrophages at various feeding concentrations of nanoparticles. (**C**) Cellular uptake rate of nanoparticles as a function of feeding nanoparticle concentrations. The data were fitted to a hyperbolic saturation model: the best-fit equation (red dashed line) is represented as *y = y_max_* × *x/(a + x*)], where, *y_max_* = 7.0 × 10^4^ represents the maximal uptake rate and *a* = 7.7 × 10^11^ is the uptake constant (nanoparticle concentration at which *y* = 0.5 *y_max_*). The shaded area depicts the 90% prediction interval. (**D**) Relationship between the percentage of internalized nanoparticles at 4 hours and the initial amount of nanoparticles added. The data are expressed as mean ± SD, with n = 4.

Next, the uptake kinetics of SiNP-R6G were measured at various treating concentrations, ranging from 6.0 × 10^10^ to 9.1 × 10^11^ nanoparticles per mL. Figure 6B demonstrates an almost linear accumulation of nanoparticles per cell over time, suggesting that the cellular uptake of nanoparticles follows apparent zero-order kinetics. The cellular uptake rate, calculated per hour and per cell, was determined by applying a linear regression to the kinetic profiles shown in Figure 6B. The resulting uptake rates (*V*) were plotted on the y-axis against the added nanoparticle concentrations (*C*) on the x-axis (Figure 6C). A hyperbolic, saturable function was fitted to the rate data, yielding the best-fit equation:

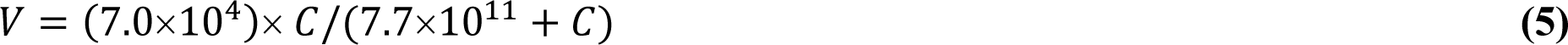

The fitting analysis revealed that the maximum uptake rate (*V_max_*) per macrophage is 7.0 × 10^4^ nanoparticles per hour, with a half-saturation concentration of 7.7 × 10^11^ nanoparticles per mL. The nonlinearity in uptake kinetics is further demonstrated in Figure 6D, which illustrates that the percentage uptake (relative to the added concentration) at the end of the incubation period (i.e. 4 hours) decreases as the concentration increases.

To assess the endocytosis capability of macrophages, we conducted calculations to estimate the total volume of intracellular nanoparticles relative to the average macrophage volume (Table 2). The numbers of particles within the cells were determined by incubating macrophages with various number concentrations of SiNP-R6G for 4 hours. The estimated maximum capacity of nanoparticle uptake at 4 hours was calculated using Equation 5. The average diameter of the nanoparticles, determined by NTA, was found to be 0.114 μm. The total volumes of nanoparticles (V*_np_*) were then calculated by multiplying the sphere volume with the estimated numbers of nanoparticles present within the cells.

**Table 2.**
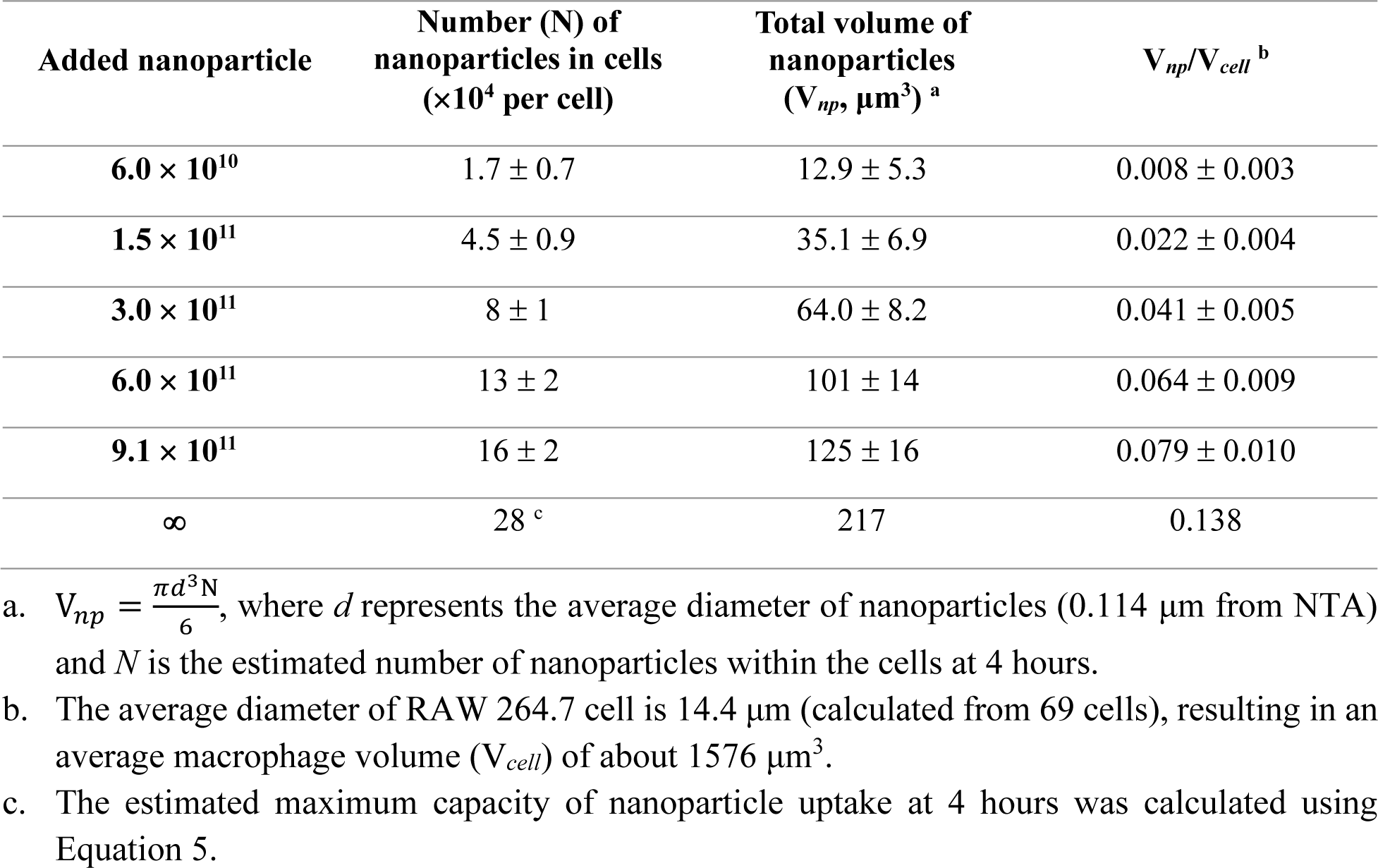
Estimated Total Volume of Intracellular Nanoparticles Relative to Average Macrophage Volume

| Added nanoparticle | Number (N) of nanoparticles in cells ($\times 10^4$ per cell) | Total volume of nanoparticles ($V_{np}$, $\mu\text{m}^3$)<sup>a</sup> | $V_{np}/V_{cell}$<sup>b</sup> |
| --- | --- | --- | --- |
| $6.0 \times 10^{10}$ | $1.7 \pm 0.7$ | $12.9 \pm 5.3$ | $0.008 \pm 0.003$ |
| $1.5 \times 10^{11}$ | $4.5 \pm 0.9$ | $35.1 \pm 6.9$ | $0.022 \pm 0.004$ |
| $3.0 \times 10^{11}$ | $8 \pm 1$ | $64.0 \pm 8.2$ | $0.041 \pm 0.005$ |
| $6.0 \times 10^{11}$ | $13 \pm 2$ | $101 \pm 14$ | $0.064 \pm 0.009$ |
| $9.1 \times 10^{11}$ | $16 \pm 2$ | $125 \pm 16$ | $0.079 \pm 0.010$ |
| ∞ | $28^c$ | 217 | 0.138 |

- a. $V_{np} = \frac{\pi d^3 N}{6}$ , where $d$ represents the average diameter of nanoparticles (0.114 $\mu\text{m}$ from NTA) and $N$ is the estimated number of nanoparticles within the cells at 4 hours. - b. The average diameter of RAW 264.7 cell is 14.4 $\mu\text{m}$ (calculated from 69 cells), resulting in an average macrophage volume ( $V_{cell}$ ) of about 1576 $\mu\text{m}^3$ . - c. The estimated maximum capacity of nanoparticle uptake at 4 hours was calculated using Equation 5.

The average diameter of RAW 264.7 cells, calculated from 69 cells, is approximately 14.4 μm, resulting in an average macrophage volume (V*_cell_*) of approximately 1576 μm^3^. The maximum volume ratio of nanoparticles to macrophages is approximately 13.8%. Previous studies have suggested that a high particle occupation of macrophage volume (i.e., > 6%) may lead to a diminished clearance rate of particles by macrophage, which could potentially benefit drug delivery systems[46, 47].

### 3.4 Visualization of SiNP-R6G Cellular Uptake

We conducted microscopic imaging studies to visually confirm the presence of nanoparticles during the cellular uptake process. Figure 7 shows the images obtained from confocal laser scanning microscopy (CLSM), providing visual evidence of the intracellular presence of SiNP-R6G within macrophages. Figure 8 further reinforces the observation of slow uptake kinetics of nanoparticles as observed in the quantitative analysis (Figure 6A & 6B). Additionally, Figure 9 demonstrates that the intracellular fluorescent signal of nanoparticles partially overlaps with that of the lysosomal tracker, indicating a distribution of nanoparticles within the lysosomes.

**Figure 7.**
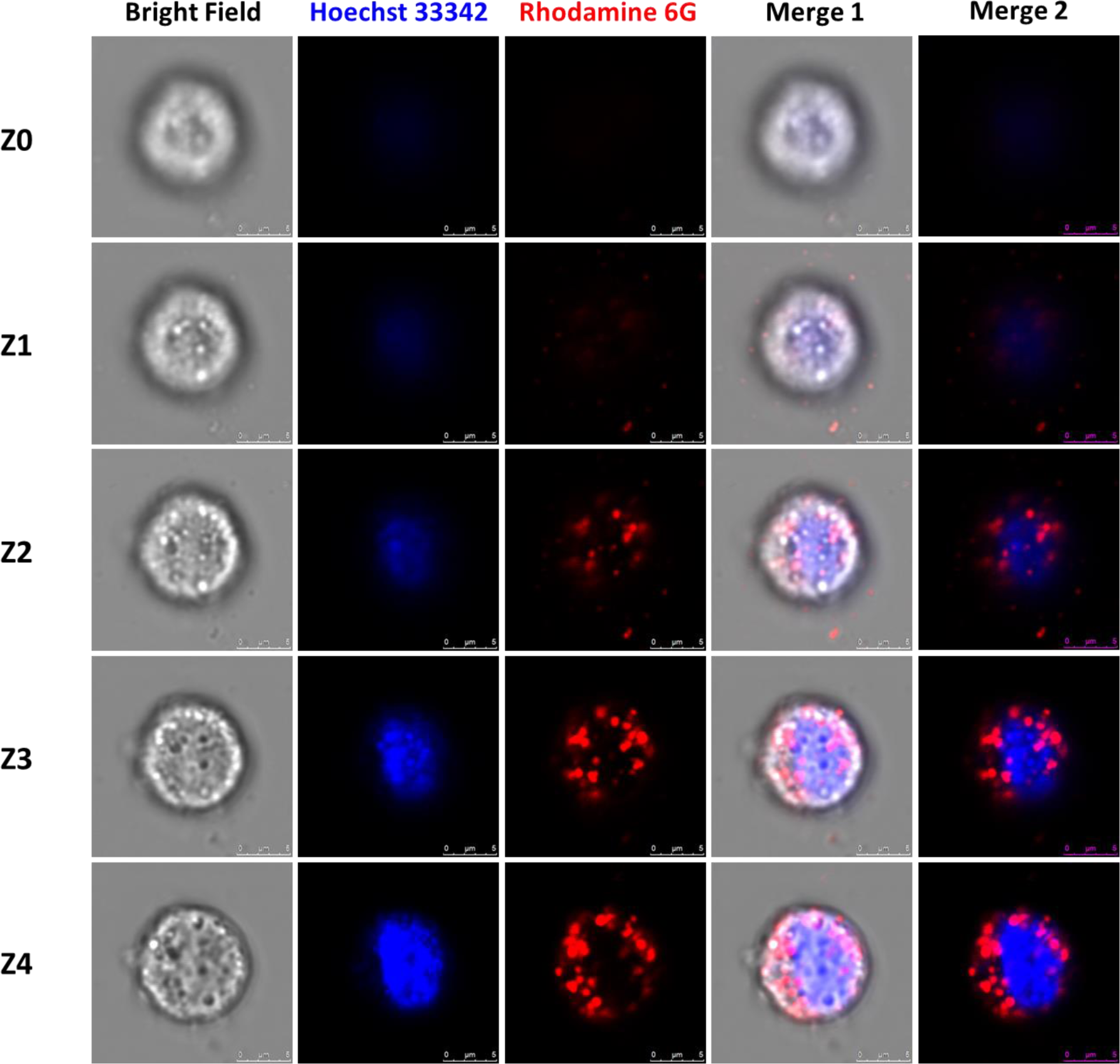
Visualization of SiNP-R6G Internalization in Macrophages Using Confocal Laser Scanning Microscopy (CLSM). RAW 264.7 cells were treated with SiNP-R6G for 4 hours. The nucleus was stained using Hoechst 33342 (blue). Five z-stack images (Z0 to Z4) of a single cell were acquired, with each image capturing a different focal depth, Z0 represents the nucleus image with the minimum contour (step size = 0.5 μm; scale bar = 5 μm). Merge 1 shows the overlap of bright field and fluorescent images, while Merge 2 displays the overlap of fluorescent images only.

**Figure 8.**
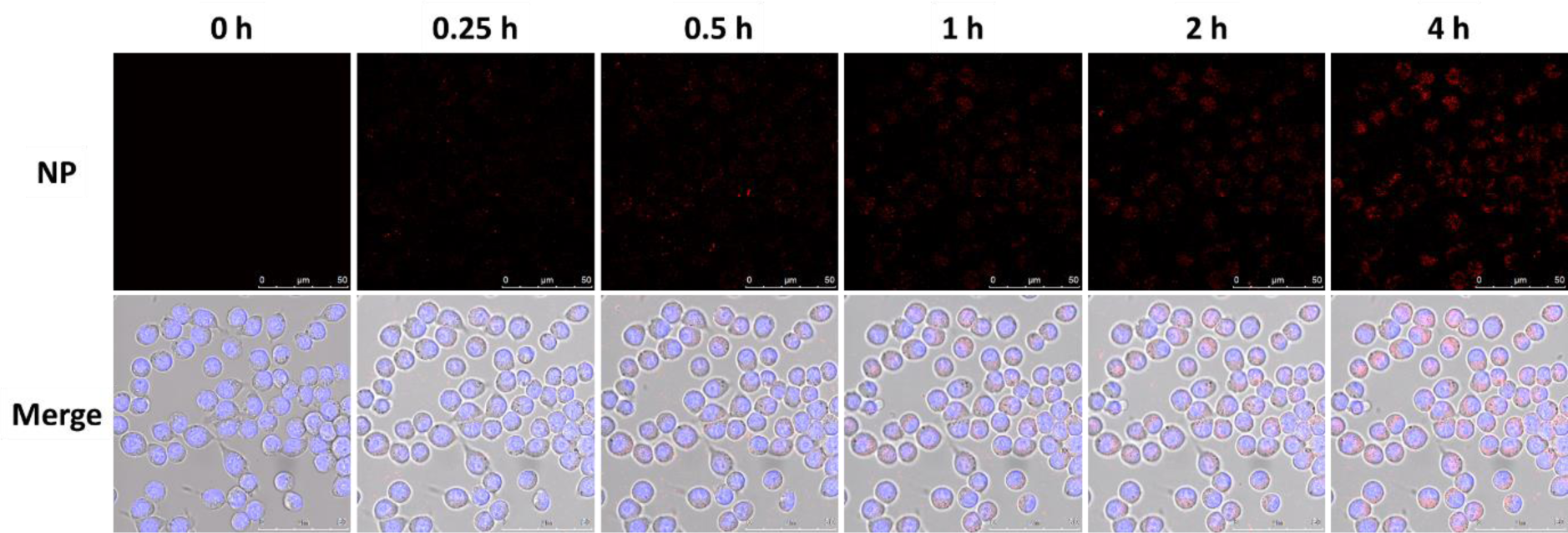
Time-lapse Fluorescent Imaging of SiNP-R6G Cellular Uptake (Red Fluorescence). CLSM images of RAW 264.7 cells were captured at various time points: before addition of SiNP-R6G (0 h) and at 0.25, 0.5, 1, 2, and 4 hours post-treatment. The lower panel displays merged images showing the blue stained nucleus and red fluorescent nanoparticles. Scale bar = 50 μm.

**Figure 9.**
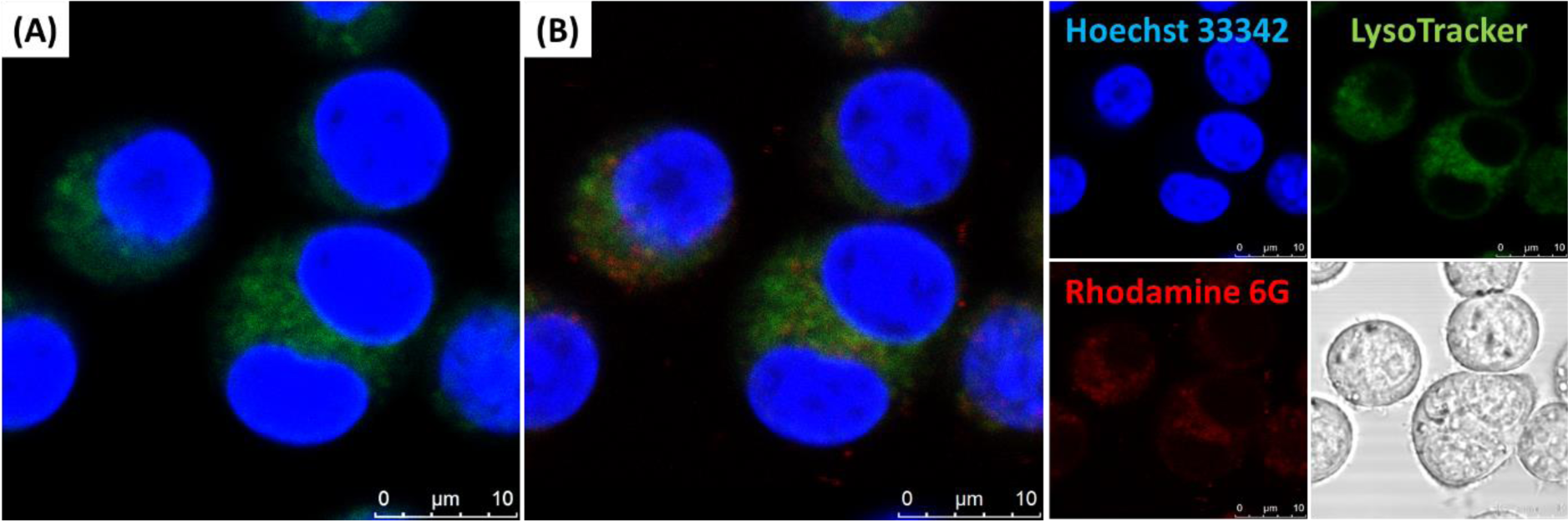
Visualization of Intracellular Localization of SiNP-R6G using CLSM. (A) Merged images of RAW 264.7 cells prior to addition of SiNP-R6G. (**B**) Merged and separate fluorescent images of RAW 264.7 cells after a 4-hour incubation with SiNP-R6G. Color representation: blue for nucleus staining (Hoechst 33342), green for lysosomal staining (LysoTracker^TM^ Deep Red), red for nanoparticles. The bright-field image of nanoparticle-treated cells is also included. Scale bar = 10 μm.

Transmission electron microscopy was employed to directly visualize intracellular nanoparticles. Figure 10 shows the accumulation of nanoparticles within numerous intracellular vesicles of varying sizes and quantities. The number of nanoparticles enclosed within each vesicle was counted, and a frequency distribution was plotted (Figure 10C). In the cross-sectioned image presented in Figure 10A, 33 vesicles were identified, with each vesicle containing an average of 24 ± 11 nanoparticles. The total number of nanoparticles in the image amounted to 319, with an average diameter of 97 ± 44 nm (measured using ImageJ). Notably, the size distribution of the internalized nanoparticles closely resembled that of the pristine SiNP-R6G particles obtained from TEM analysis (which exhibited a size distribution of 95 ± 34 nm). This outcome indicates that the physical characteristics of SiNP-R6G nanoparticles remain relatively intact following cellular internalization.

**Figure 10.**
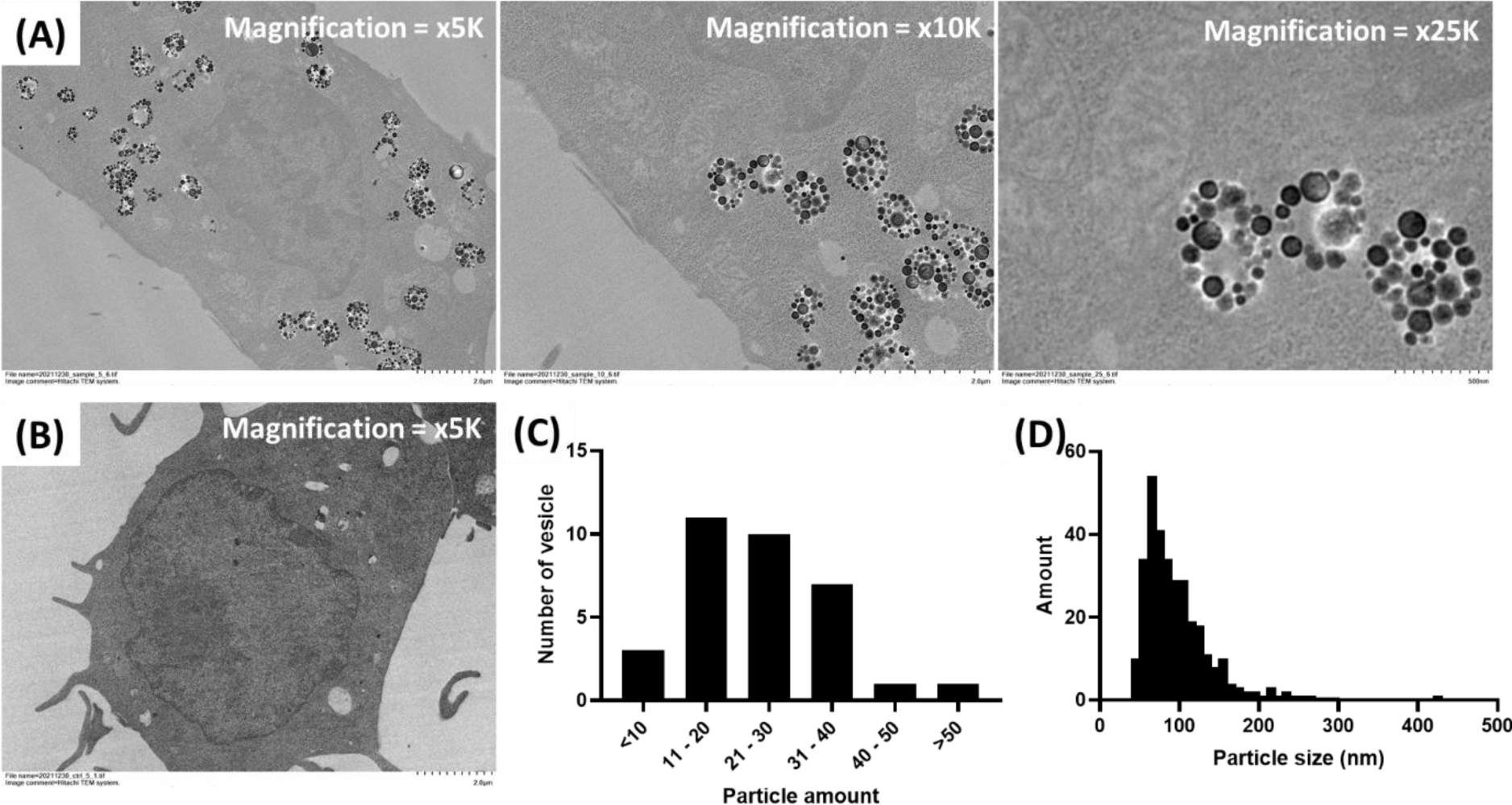
Localization of Nanoparticles within Intracellular Vesicles in Macrophages Observed by TEM. (**A**) TEM images of a macrophage at increasing magnifications (from left to right) captured after a 4-hour incubation with SiNP-R6G. Scale bar = 2 μm (left and central images) or 0.5 μm (right). (**B**) TEM image of a control macrophage without nanoparticle treatment. Scale bar = 2 μm. (**C**) Quantification of the number of particles within each intracellular vesicle in image A (x5K magnification). The frequency histogram is displayed. Number of vesicles counted = 33. (**D**) Size distribution of all nanoparticles observed in image A (left). Number of nanoparticles counted = 319.

### 3.5 Clearance of Internalized SiNP-R6G from Macrophages

To investigate the retention duration of internalized SiNP-R6G in macrophages, the cells were initially treated with SiNP-R6G for 4 hours to allow internalization. Subsequently, the remaining intracellular fluorescence was measured over time in a nanoparticle-free medium, with free R6G molecules included for comparison. Figure 11A illustrates a slow and minor decrease in cellular fluorescence for SiNP-R6G within the first 60 minutes, whereas internalized free R6G exhibited a significantly higher rate and extent of leaving the cells, reaching equilibrium rapidly within 60 minutes. Remarkably, SiNP-R6G appeared to persist within macrophages during an extended observation period of up to 2 days (Figure 11B). Additionally, a complementary CLSM study was conducted to demonstrate the prolong residence of SiNP-R6G in macrophages. As depicted in Figure 12 (A, B, E), the fluorescence images of cells containing SiNP-R6G nanoparticles were nearly identical between the initial state (A) and the 24-hour state (B). In comparison, the cellular fluorescence of free R6G substantially diminished after 1 hour (C, D, E).

**Figure 11.**
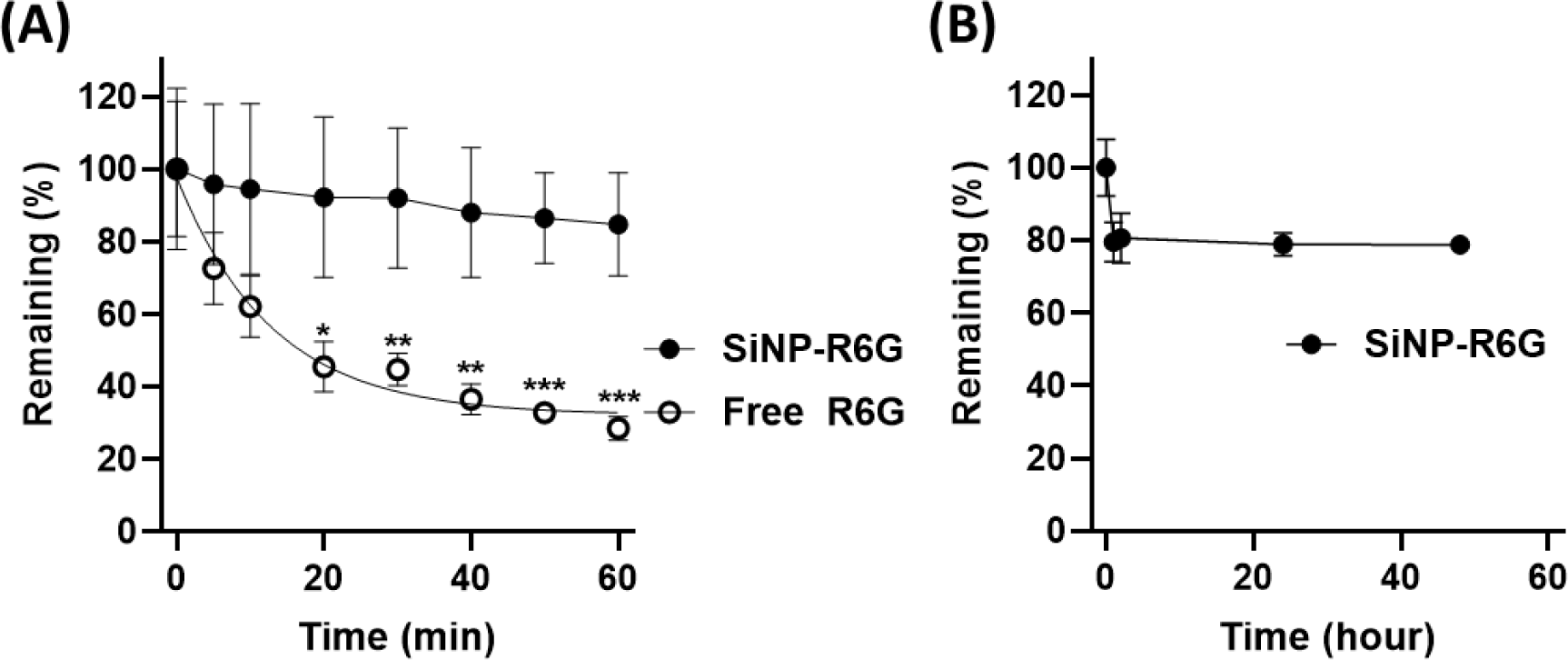
Elimination Kinetics of Internalized Free R6G and SiNP-R6G from Macrophages. (**A**) Comparison of free 6G and SiNP-R6G within 1 hour post-uptake (loading [R6G] equivalent to 1 μM). (**B**) Extended measurement over 48 hours for SiNP-R6G (loading [R6G] equivalent to 10 μM). Macrophage cells were treated with either free R6G or SiNP-R6G for 4 hours. After removing the loading solution and washing with PBS three times, the fluorescence intensity (expressed as % remaining) of the cells was measured over time. Data are presented as mean ± SD, n = 3 or 4. * *p* < 0.05, ** *p* < 0.01, *** *p* < 0.001.

**Figure 12.**
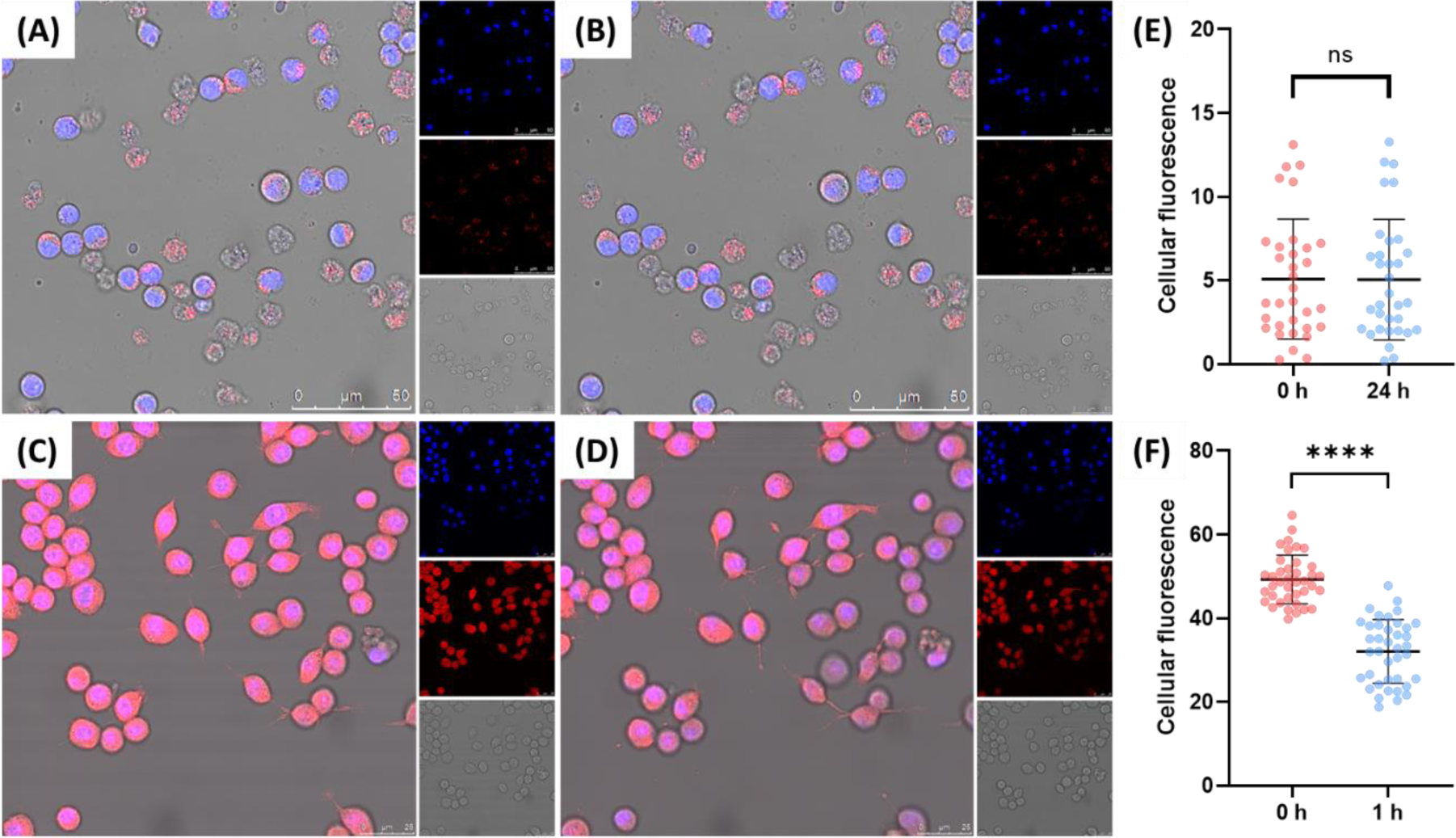
CLSM Visualization of SiNP-R6G (A, B, E) or free R6G (C, D, F) Retention in Macrophages. RAW 264.7 cells were incubated with SiNP-R6G or free R6G for 4 hours. Subsequently, the medium was replaced with a dye-free medium. (**A**) Images of cells with SiNP-R6G taken immediately after medium replacement. (**B**) Images of the same cells with SiNP-R6G taken 24 hours later. (**C**) Images of cells with free R6G taken immediately after medium replacement. (**D**) Images of the same cells with free R6G taken 1 hour later. The nucleus was stained with Hoechst 33342 (blue). Each panel shows the merged image along with individual images. Scale bar = 50 μm or 25 μm. ImageJ analysis of cellular fluorescence intensity for (**E**) cells with SiNP-R6G (number of cells = 33 in images A & B) and for (**F**) cells with free R6G (number of cells = 39 in images C &D). **** *p* < 0.0001 (paired t-test).

## 4. Discussion

Fluorescence labeling of nanoparticles is a convenient approach for tracing nanoparticle uptake and trafficking in cells and tissues. While the method allows for fluorescence intensity-based comparisons, whether in microscopic images or through a cell sorter, associating the intensity information with the exact quantity of nanoparticles in the sample poses a challenge. In this study, we aim to enhance the capabilities of fluorescence based measurements by establishing a quantitative relationship between fluorometric measurements of fluorescent nanoparticles and nanoparticle tracking analysis (NTA). The primary objective is to quantify the kinetics of nanoparticle uptake in a model murine macrophage, RAW 264.7. Ultimately, we seek to answer questions related to nanoparticle uptake kinetics, such as: What is the macrophage’s internalization rate of nanoparticles in terms of particles per hour? Or, what is the maximum particle capacity a macrophage can uptake? We first demonstrate the essential calibration procedure required for the proposed quantitative approach. After successfully validating the method, we effectively convert the fluorometric kinetic data into countable particle numbers within cells. To corroborate the quantitative data, we provide clear visual evidence of nanoparticle crowding within intracellular vesicles, an intriguing phenomenon that has not received much attention previously.

Our study revealed that the initial internalization of organosilica nanoparticles (with a diameter of approximately 100 nm) in macrophages follows a slow and constant zero-order kinetics (Figure 6). The constant-rate results support the notion that a carrier/vehicle system in involved in delivering extracellular particle cargos into the cells. The hypothesis was confirmed by TEM images, which showed intracellular vesicular compartments densely packed with nanoparticles (Figure 10). Thus, the nanoparticles are taken up as a group rather than individually. Importantly, each individual nanoparticle within the intracellular vesicles was clearly visible and quantifiable. It is rare, if not unprecedented, to observe an intracellular vehicle capable of accommodating nearly 50 nanoparticles at maximum capacity (Figure 10).

Our method offers the potential to address inquiries regarding the rate and extent of nanoparticle internalization and residence within the model macrophage. The observed rate of internalization ranges from 4.0 × 10^3^ to 3.7 × 10^4^ nanoparticles per hour, depending on the concentration of loaded nanoparticles (Figure 6C). The estimated maximum uptake rate is 7.0 × 10^4^ nanoparticles per hour, with a half-saturation concentration of approximately 0.8 trillion nanoparticles per mL. Importantly, our findings align with a recent study demonstrating the high capacity of macrophages to take up pegylated gold nanoparticles [41]. Furthermore, it is noteworthy that the internalized nanoparticles persist within the macrophages for an extended period (Figure 11B), without significantly affecting cell viability (Figure 4A). These findings emphasize the potential of utilizing macrophages as the “Trajan Horse”, carrying a substantial load of nanoparticle “soldiers” to the targeted therapeutic site [36–40].

## 5. Conclusions

Our study represents a breakthrough in the field of fluorescent nanoparticle tracking, particularly in the context of quantifying nanoparticle uptake kinetics. Through the establishment of a quantitative relationship between fluorometric measurements and nanoparticle tracking analysis, we were able to accurately convert fluorometric kinetic data into numerical particle counts within cells. This achievement effectively overcomes the limitations associated with comparisons based solely on fluorescence intensity. Importantly, our findings shed light on the previously underappreciated phenomenon of nanoparticle crowding within intracellular vesicles. This discovery could potentially stimulate further investigation into the implications and consequences of intracellular nanoparticle crowding, which has the capacity to influence the design and optimization of nanoparticle-based therapies and drug delivery systems.

## Author Information

Corresponding Author Shih-Jiuan Chiu − School of Pharmacy, Taipei Medical University, Taipei 110, Taiwan; orcid.org/0000-0001-9952-4055; Teh-Min Hu – Department of Pharmacy, National Yang-Ming Chiao Tung University, Taipei 112, Taiwan; orcid.org/0000-0001-6350-3180;

## Acknowledgements

This study received support from the National Science and Technology Council, Taiwan. We gratefully acknowledge Yen-Ju Chan, Ph.D. for providing assistance with confocal laser scanning microscopy, and Mr. Clement Lee, Deputy Director of the Core Facility Center in TMU, for their valuable contribution in preparing TEM samples.

